# Mechanical and tactile incompatibilities cause reproductive isolation between two young damselfly species

**DOI:** 10.1101/142828

**Authors:** Alexandra A. Barnard, Ola M. Fincke, Mark A. McPeek, John P. Masly

## Abstract

External male reproductive structures have received considerable attention as an early-acting cause of reproductive isolation (RI), because the morphology of these structures often evolves rapidly between populations. This presents the potential for mechanical incompatibilities with heterospecific female structures during mating and could thus prevent interbreeding between nascent species. Although such mechanical incompatibilities have received little empirical support as a common cause of RI, the potential for mismatch of reproductive structures to cause RI due to incompatible species-specific tactile cues has not been tested. We tested the importance of mechanical and tactile incompatibilities in RI between *Enallagma anna* and *E. carunculatum,* two damselfly species that diverged within the past ~250,000 years and currently hybridize in a sympatric region. We quantified 19 prezygotic and postzygotic RI barriers using both naturally occurring and lab-reared damselflies. We found incomplete mechanical isolation between the two pure species and between hybrid males and pure species females. Interestingly, where mechanical isolation was incomplete, females showed greater resistance and refusal to mate with hybrid or heterospecific males compared to conspecific males, which suggests that tactile incompatibilities involving male reproductive structures can influence female mating decisions and form a strong barrier to gene flow in early stages of speciation.

## Introduction

Understanding speciation requires identifying how reproductive isolation (RI) is initiated and maintained in the early stages of population divergence (Coyne and Orr 2004; Butlin et al. 2012). Over the past century, speciation researchers have used a variety of experimental and comparative approaches to identify which barriers appear most important in causing RI early in the speciation process. These efforts have revealed that sexual isolation and ecological divergence tend to evolve earlier than hybrid sterility and inviability in both plants (*e.g.*, Grant 1992; Ramsey et al. 2003; Husband and Sabara 2004; Kay 2006) and in animals (*e.g.*, McMillan et al. 1997; Price and Bouvier 2002; Mendelson and Wallis 2003; Dopman et al. 2010; Sánchez-Guillén et al. 2012; Williams and Mendelson 2014; Castillo et al. 2015). Prezygotic isolation also typically evolves faster in sympatry than in allopatry, and hybrid sterility typically evolves faster than hybrid inviability (Coyne and Orr 1997; Presgraves 2002; Price and Bouvier 2002; Russell 2003). Identifying the traits that diverge to cause RI underlying these broad patterns is a major goal of speciation research.

One set of traits that has received much attention because of their rapid rates of evolutionary change is external reproductive structures. In internally fertilizing animals, male intromittent genitalia are among the fastest-evolving external morphological traits, and genital morphological variation can affect reproductive fitness within species (Eberhard 1985; Otronen 1998; Danielsson and Askenmo 1999; House and Simmons 2003; Rodriguez et al. 2004; Bertin and Fairbairn 2005; Simmons et al. 2009). Likewise, non-intromittent contact or grasping structures often show similar patterns of rapid, divergent evolution, and divergence in these structures can also affect reproductive success within species (Arnqvist 1989; Bergsten et al. 2001; Wojcieszek and Simmons 2012).

Rapid divergence of reproductive structures between populations has been hypothesized to cause RI via two different mechanisms. The first is mechanical incompatibility (Dufour 1844), in which structural incompatibilities between male and female genitalia of different species prevent successful copulation and reproduction. Mechanical incompatibilities have been documented in some animal species pairs (Jordan 1896; Standfuss 1896; Federley 1932; Schick 1965; Paulson 1974; Sota and Kubota 1998; Tanabe and Sota 2008; Kamimura and Mitsumoto 2012; Sánchez-Guillén et al. 2012; Wojcieszek and Simmons 2013; Sánchez-Guillén et al. 2014; Anderson and Langerhans 2015), although this mechanism of RI has not received broad support as a common mechanism of RI between young species (Shapiro and Porter 1989; Masly 2012; Simmons 2014).

The second proposed mechanism is tactile incompatibility (de Wilde 1964; Eberhard 1992), in which mismatch between male and female genitalia of different species prevents or reduces the success of mating and reproduction because one or both sexes fail to stimulate the other in the proper species-specific manner. The essence of this idea is that female reproductive decisions are based on the pattern of tactile stimuli transmitted by the male, and improper stimulation can result in female refusal to mate, early termination of mating, or lowered postcopulatory fitness, including reduced reproductive fitness in hybrid offspring (Eberhard 2010). Tactile isolation likely operates in a similar manner as other sensory modalities involved in mate choice and species recognition such as auditory or chemical signals, in which quantitative variation exists in male traits and female preferences (Ryan and Wilczynski 1991; Shaw 1996; Tregenza and Wedell 1997; Singer 1998; Johansson and Jones 2007). If females discriminate among the mating structures of conspecific mates, female discrimination against heterospecific males can arise as a byproduct of sexual selection within species (reviewed in Panhuis et al. 2001; Turelli et al. 2001; Simmons 2014). Thus, any mismatch between male morphology and female response to stimulation from a particular morphology could result in reduced reproductive success when females mate with a heterospecific male or an interspecific hybrid male. The importance of tactile incompatibilities remains unknown, although there is good reason to expect that these incompatibilities may occur frequently (Simmons 2014), and therefore have the potential to play a significant role in the evolution of RI.

Because identifying the effects of tactile incompatibilities requires carefully quantifying mating behavior and physiology, these incompatibilities have often been overlooked in tests of RI involving divergence of reproductive structures (Masly 2012). Nonetheless, some evidence for tactile incompatibility in the absence of mechanical incompatibilities exists in butterflies (Lorkovic 1953, 1958), scarab beetles (Eberhard 1992), *Drosophila* (Coyne 1993; Price et al. 2001; Frazee and Masly 2015 – but see LeVasseur-Viens et al. 2015), and sepsid flies (Eberhard 2001). Notably, damselflies (Odonata, suborder Zygoptera) are often touted as a prime example of the importance of both mechanical and tactile incompatibilities in RI among closely related species. The potential for either mechanism to cause RI has been particularly well described in the families Lestidae and Coenagrionidae, whose males do not engage in premating courtship or visual displays (Williamson 1906; Krieger and Krieger-Loibl 1958; Loibl 1958; Paulson 1974; Tennessen 1975; Robertson and Paterson 1982; Hilton 1983; Battin 1993; Sánchez-Guillén et al. 2012; Sánchez-Guillén et al. 2014). Male damselflies have two sets of paired grasping organs at the end of their abdomen (Fig 1). A male initiates the mating sequence by grasping the female’s thorax with these appendages to form the “tandem” position. The species-specific male appendages and female thoracic structures engage such that structural mismatch appears to prevent many heterospecific tandems from forming. Mechanical isolation appears to be a major cause of RI in *Ischnura* (Krieger and Krieger-Loibl 1958; Sánchez-Guillén et al. 2014). For *Ischnura* species pairs with incomplete mechanical isolation, tactile isolation has been suggested to contribute to RI (Sánchez-Guillén et al. 2012; Wellenreuther and Sánchez-Guillén 2016), although this idea has not been tested quantitatively.

**Fig 1.**
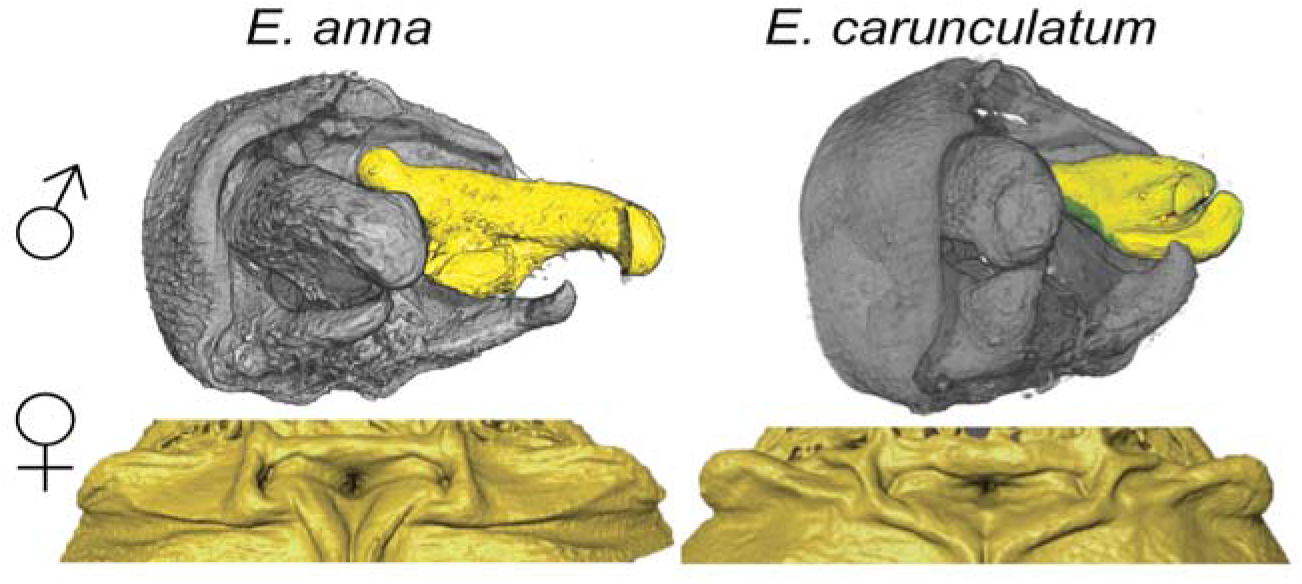
Male grasping appendages and female mesostigmal plate morphology. The right cercus on each male is shaded yellow.

Mechanical isolation also appears to prevent many heterospecific tandems in *Enallagma,* the most speciose North American genus (Paulson 1974; Miller and Fincke 2004; Fincke et al. 2007). Divergence in reproductive structure morphology is associated with a relatively recent *Enallagma* radiation (250,000-15,000 years ago; McPeek et al. 2008). Importantly, the rapid morphological diversification was not accompanied by marked ecological divergence among many *Enallagma* species (Siepielski et al. 2010). Although male cerci (superior terminal appendages) and female thoracic plates show a pattern of correlated evolution within *Enallagma* species (McPeek et al. 2009), species-specific divergence in these structures does not always cause strong mechanical incompatibilities, and interspecific tandems are occasionally observed (Paulson 1974; Tennessen 1975; Bick and Bick 1981; Forbes 1991; Miller and Fincke 2004; Fincke et al. 2007). After tandem formation, female *Enallagma* control whether or not copulation occurs and they typically refuse to mate with heterospecifics or males whose cercus morphology has been manipulated (Robertson and Paterson 1982). *Enallagma* mesostigmal plates contain mechanoreceptors in species-specific locations that appear to be contacted by the male cerci during tandem, which may allow female assessment of a male’s cercus morphology (Robertson and Paterson 1982).

Although prezygotic isolating barriers appear to evolve earlier than postzygotic barriers in damselflies (Sánchez-Guillén et al. 2012; Sánchez-Guillén et al. 2014), the relative importance of mechanical and sensory mechanisms of prezygotic RI remains unclear for two reasons. First, it can be difficult to distinguish between mechanical and tactile mechanisms experimentally: if a male-female pair fails to form a tandem, it is often unclear whether the incompatibility is purely mechanical or whether it involves tactile or behavioral cues that cause one sex to reject the other (Tennessen 1975; Robertson and Paterson 1982; Shapiro and Porter 1989). Second, mechanical isolation or male-female “fit” is not always defined in a way that makes quantifying variation in these phenotypes straightforward (Masly 2012). This lack of clarity over what constitutes mechanical incompatibility has led to conflation of mechanical RI (*i.e.*, failure of male and female parts to engage) in damselflies with mechanisms that might be better described as tactile (Tennessen 1982).

Distinguishing mechanical from tactile mechanisms requires performing detailed mating observations among males and females that possess interspecific variation in reproductive structures and identifying specific features of reproductive morphology that prevent mating or reduce mating success using high-resolution phenotypic data. Here, we take advantage of a large collection of naturally occurring interspecific hybrids and lab-generated hybrids to test the hypothesis that divergence in reproductive structural morphology causes RI at the early stages of speciation in damselflies. We measure 19 potential pre- and postzygotic isolating barriers between *Enallagma anna* and *E. carunculatum,* two species that diverged from a common ancestor sometime in the last ~250,000 generations (McPeek et al. 2008; Callahan and McPeek 2016) and co-occur over much of the western United States (Westfall and May 2006). Both species have identical ecologies and overall morphologies (Turgeon et al. 2005; McPeek et al. 2009), but display conspicuous differences in the size and shape of the male cerci and female mesostigmal plates (Fig 1). We quantify variation in male and female reproductive structure morphologies, distinguish mechanical and tactile premating incompatibilities, estimate the cumulative strengths of multiple reproductive barriers, and independently test predictions of mechanical and tactile isolation hypotheses (Richards and Robson 1926; Shapiro and Porter 1989). If mechanical incompatibilities occur, male *E. anna* × *E. carunculatum* hybrids that possess intermediate cercus morphologies will have less success at forming tandems compared to conspecific males. If tactile incompatibilities occur, males will be able to achieve tandem regardless of their cercus morphology, but females will refuse to mate with males whose morphologies deviate significantly from the conspecific mean phenotype.

## Materials and Methods

Damselfly cerci and mesostigmal plates are non-intromittent sexual structures that are not directly involved in the transfer of gametes from male to female. However, terminal appendages of male insects and the female structures they contact during mating are often referred to as secondary genital structures. We thus include them as genital traits, consistent with previous definitions (Eberhard 1985; Arnqvist and Rowe 2005; Eberhard 2010; Simmons 2014; Brennan 2016) and refer to them generally as “genitalia” in the presentation of our results.

### Natural population sampling

We studied wild populations of *E. anna* and *E. carunculatum* in July and August 2013 at a site on the Whitefish River (Montana, U.S.A.; 48°22'15"N 114°18'09"W), where putative interspecific hybrids have been reported (Miller and Ivie 1995; Westfall and May 2006). To estimate relative frequencies of each species, we collected solitary males and tandem/copulating male-female pairs during peak activity between 1030-1600 hr. We initially assigned species identity after inspecting cercus and mesostigmal plate morphology with a hand lens or dissecting microscope, respectively. Males and females with morphologies that appeared intermediate were initially designated as hybrids. We reassessed these assignments in the lab after 3-D morphometric analysis (see below). We calculated the proportions of *E. anna, E. carunculatum* and hybrid males from all sampling bouts and used these male frequencies to estimate the expected frequencies of each type of male-female pair under random mating.

We attempted to cross virgin *E. anna* and *E. carunculatum* to measure postzygotic RI between pure species, but we did not obtain heterospecific copulations in either cross direction. Instead, we established laboratory populations of hybrids and parental species by collecting eggs from mated pairs captured in the field. Mated females oviposited on moist filter paper, which was kept submerged in 2-4 cm of water until larvae hatched. We obtained embryos from 24 *E. anna* pure species crosses, 32 *E. carunculatum* pure species crosses, and 8 mixed crosses: 1 *E. carunculatum* female × *E. anna* male, 1 *E. anna* female × *E. carunculatum* male, 1 *E. carunculatum* female × hybrid male, and 5 hybrid female × *E. anna* male (“hybrid” refers to damselflies with intermediate cercus or mesostigmal plate morphologies). After sampling, egg collection, and behavioral observation, we stored adult damselflies in 95% ethanol for subsequent morphometric analyses.

### Laboratory rearing

We transported embryos from the field site to the University of Oklahoma Aquatic Research Facility where the larvae hatched and were reared to adulthood in individual 140 ml cups. The larvae were provided with *Artemia, Daphnia,* or *Lumbriculus* as food sources and experienced a natural photoperiod and daily water temperatures that averaged 20.0 + 0.19 °C. We housed adults in mesh cages (30.5 cm^3^; BioQuip), segregated by sex until sexual maturity and provided with adult *Drosophila* as a food source *ad libitum*. We used lab-reared virgin adults to quantify prezygotic barriers, plus additional postzygotic barriers that we could not measure in the field. We mated 24 adult pairs from this first lab generation: 11 *E. anna,* 2 *E. anna* female × hybrid male, 2 *E. carunculatum* female × hybrid male, 6 hybrid female × *E. anna* male and 4 hybrid female × hybrid male. Embryos from the second lab generation contributed fecundity, fertility, and hatch rate data but were not raised to adulthood due to difficulties with rearing them. Mated adults were stored in 95% ethanol after mating (males) or after oviposition (females). Unmated damselflies were maintained to calculate captive lifespan, then preserved in 95% ethanol.

### Morphometric analysis

We photographed ethanol-preserved adults using a Nikon D5100 camera (16.2 MP; Nikon Corporation, Tokyo, Japan) and measured abdomen length (abdominal segments 1-10, excluding terminal appendages) as a proxy for body size using ImageJ (Abramoff et al. 2004) for 175 males and 171 females. To reduce measurement error, we measured each abdomen twice, then used the mean length in subsequent analyses after confirming that repeatability was high for the separate measurements (*r* = 0.97). We obtained 3-D digital reconstructions of male cerci and female mesostigmal plates by scanning 140 male terminal segments and 162 female thoraces in a SkyScan 1172 micro-computed tomography scanner (Bruker microCT, Kontich, Belgium). Male structures were scanned at a voxel resolution of 2.36 or 2.53 um, and female thoraces at 2.78 or 3.88 um, and the scan data were converted to image stacks using NRecon version 1.4.4 (Bruker microCT).

To quantify cercus shape, we digitally segmented the right cercus from each male’s image stack and converted it to a solid surface object using Avizo Fire software (FEI Software; Hillsboro, Oregon) as described in McPeek et al. (2008). We measured the volume of each cercus object as a proxy for cercus size, using Avizo’s volume measurement tool. To quantify and compare their shapes, each cercus was represented by a mesh of 20,000 triangles with 10,002 vertices, each defined by distinct (*x, y, z*) coordinates (Fig S1). We placed 7 landmarks on common points on each cercus, then used these landmarks to register all digitized cerci in identical orientations within the coordinate plane. To ensure that only shape and not size was compared in the analysis, all objects were standardized to have the same centroid size. Next, we performed spherical harmonic analysis (Shen et al. 2009), which represents the shape of a closed surface in terms of the sum of 3-D sines and cosines on a sphere. We performed the analysis using 18 degrees of spherical harmonic representation, which captures relevant surface detail without introducing excess noise (Shen et al. 2009). The analysis generated 1,083 coefficients to describe the shape of each cercus, which we reduced into the primary axes of shape differentiation using principal component analysis.

Because female mesostigmal plates are relatively flat structures, we represented plate morphology using 3-D geometric morphometrics. For each female plate we assigned 11 fixed landmarks and 248 sliding semi-landmarks to the right anterior thorax of each female (Fig S2) using Landmark software (Wiley et al. 2005). We imported landmark coordinates into R and used the Geomorph package (version 2.1.7; Adams and Otarola-Castillo 2013) to assign 79 landmarks as “curve sliders” on the medial thorax and around the plate periphery, and 169 “surface sliders” evenly spaced across the plate. We obtained 3-D shape variables for these representations using general Procrustes analysis superimposition (Rohlf 1999), then obtained a smaller set of plate shape variables from the Procrustes-superimposed coordinates using principal component analysis.

### Measuring pre- and postzygotic reproductive isolating barriers

To measure the strength of RI barriers between *E. anna* and *E. carunculatum,* we quantified 19 potential pre- and postzygotic isolating mechanisms that act from the beginning of the mating sequence through an individual’s life history. Table 1 summarizes these RI measures and describes the equations used to estimate the absolute strength of each (Dopman et al. 2010).

**Table 1.** Formulas for the absolute strength of each reproductive isolating barrier measured, listed in the order in which they act during the mating sequence and subsequent life history of an individual. In the postzygotic barrier formulas, “heterospecific” includes male-female pairs composed of both pure species and any male-female pair involving at least one hybrid partner.

| Barrier | RI formula |
| --- | --- |
| Prezygotic |  |
| Visual | $1 - (\text{number heterospecific tandem attempts} / \text{conspecific tandem attempts})$ |
| Precopulatory mechanical | $1 - (\text{number heterospecific tandems} / \text{number heterospecific tandem attempts})$ |
| Tactile I (female resistance) | $1 - (\text{proportion heterospecific tandems without resistance} / \text{proportion conspecific tandems without resistance})$ |
| Tactile II (female refusal) | $1 - (\text{proportion heterospecific matings} / \text{proportion conspecific matings})$ |
| Postzygotic |  |
| Hybrid mechanical I (tandem) | $1 - (\text{number hybrid tandems} / \text{number hybrid tandem attempts})$ |
| Hybrid mechanical II (intromission) | $1 - (\text{number hybrid copulations} / \text{number hybrid intromission attempts})$ |
| Hybrid tactile I (female resistance) | $1 - (\text{proportion hybrid tandems without resistance} / \text{proportion conspecific tandems without resistance})$ |
| Hybrid tactile II (female refusal) | $1 - (\text{proportion hybrid matings} / \text{proportion conspecific matings})$ |
| Hybrid copulation duration | $1 - (\text{mean hybrid copulation duration} / \text{mean conspecific copulation duration})$ |
| Hybrid copulation interruption duration | $1 - (\text{mean conspecific copulation interruption duration} / \text{mean hybrid copulation interruption duration})$ |
| Hybrid oviposition | $1 - (\text{proportion females oviposited, hybrid matings} / \text{proportion females oviposited, conspecific matings})$ |
| Fecundity | $1 - (\text{mean number eggs, hybrid clutch} / \text{mean number eggs, conspecific clutch})$ |
| Fertility | $1 - (\text{mean fertilized eggs, hybrid clutch} / \text{mean fertilized eggs, conspecific clutch})$ |
| Egg hatching | $1 - (\text{proportion hatched eggs, heterospecific clutch} / \text{proportion hatched eggs, conspecific clutch})$ |
| Embryo development | $1 - (\text{mean days from oviposition to hybrid egg hatching, hybrids} / \text{mean days from oviposition to pure species egg hatching})$ |
| Larval maturation time | $1 - (\text{mean days from pure species hatch to adult emergence} / \text{mean days from hybrid hatch to adult emergence})$ |
| Larval survivorship | $1 - (\text{proportion hybrid larvae that reach adulthood} / (\text{proportion pure species larvae that reach adulthood}))$ |
| Adult sex ratio | $1 - (\text{hybrid sex ratio} / \text{pure species sex ratio})$ |
| Adult lifespan | $1 - (\text{hybrid lifespan} / \text{pure species lifespan})$ |

#### Mate discrimination

We measured males’ visual discrimination of potential mates by restraining individual *E. anna* and *E. carunculatum* females on wooden dowels near the water and measuring the frequencies of each type of male that attempted tandem with them. We attached live females of each species by their legs to wooden dowels using Duco cement (ITW Devcon, Glenview, IL, USA; Miller and Fincke 1999) and placed individual dowels level with surrounding vegetation within 5 m of the water’s edge at the field site. Over 20-minute intervals, we captured each male that either attempted or achieved tandem with a restrained female and assigned them to species by examining the cerci with a hand lens. Males were held in paper envelopes until the end of the observation period to prevent the possibility of a second encounter with the restrained female and were then released.

#### Mating assays

We measured several premating RI barriers using no-choice mating experiments in which females were placed in mesh cages with either heterospecific, hybrid, or conspecific males. We used both field-caught and lab-reared damselflies, and used only virgin females in each mating assay. To obtain virgins in the field, we captured newly emerged females, identified by their pale teneral coloration. We assigned species identity as described above, then housed virgin females in cages until they reached sexual maturity (~10 days post-emergence). We placed 2-5 individuals of each sex in a cage under partial shade in the grass and observed behaviors between 1000-1600 hr.

We quantified precopulatory mechanical RI by measuring the frequency of tandem attempts in which the male was unable to securely grasp a female for longer than five seconds. A secure hold was confirmed by observing the male flying while engaged with the female, or attempting to fly without losing contact while the female remained perched. We measured copulatory mechanical RI as the proportion of copulation attempts in which the male and female failed to achieve genital coupling. This estimates mechanical incompatibility between male grasping appendages and female thoracic plates and excludes the possibility of male loss of interest, because males were often observed repeatedly attempting tandem on the same female despite being unable to grasp her.

We quantified two types of precopulatory tactile incompatibilities using pairs that formed tandems. First, we recorded whether each female showed resistance behaviors during tandem (e.g., head shaking, wing flapping, dorsal abdominal extension, or body repositioning) (Tennessen 1975; Xu and Fincke 2011). Second, we recorded whether females in tandem cooperated in copulation or refused to mate.

#### Postzygotic isolation

We quantified several postmating RI barriers using the progeny from interspecific crosses, beginning at copulation (Table 1). We measured oviposition success as the proportion of females from each cross type that oviposited. When females failed to oviposit within three days post-mating, we checked their male partners for motile sperm by anesthetizing them with CO_2_, immediately dissecting out the seminal vesicle, gently squashing it under a coverslip, and examining the contents under a Zeiss Axio Imager 2 stereomicroscope (100× total magnification). We dissected females that failed to oviposit to check the oviduct for mature eggs and the bursa copulatrix for sperm.

We calculated fecundity by counting all eggs laid by each mated female within three days of mating. The date of egg hatch was recorded as the first day that larvae were observed. We calculated the proportion of eggs that hatched from each clutch by counting the number of unhatched eggs that remained in the filter paper seven days after first hatch. We calculated fertility of lab-reared matings by counting the number of fertilized eggs, as indicated by a dark spot that develops on the apical end of the egg (Corbet 1999). We calculated embryo development timing as days from oviposition to egg hatch, larval maturation as days from egg hatch to adult emergence, larval survivorship as the proportion of hatched larvae that emerged as to adults, plus adult sex ratio and total adult lifespan.

#### Strength of RI barriers

We estimated the absolute strength of each individual barrier using the following general equation (Ramsey et al. 2003; Dopman et al. 2010):

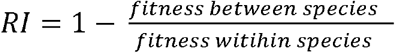

This equation yields a value between -1 and 1 in which 0 indicates no barrier to gene flow, 1 indicates full RI, and negative values indicate a hybrid fitness advantage. We estimated each RI barrier’s sequential strength (*SS*) based on its absolute strength (*AS*) and the absolute strengths of all preceding barriers, as described in (Dopman et al. 2010):

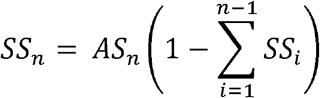

We calculated total RI (*T*) between *E. anna* and *E. carunculatum* as the sum of the sequential strengths of all barriers, then calculated each barrier’s relative contribution to total RI (*SS_n_* / *T*) (Dopman et al. 2010; Table S1).

### Statistical analyses

We compared males’ sexual approaches toward con- and heterospecific females, observed vs. expected frequency of heterospecific pairs, and adult sex ratios using binomial tests. We compared presence or absence of female resistance behaviors, female mating refusal or cooperation, frequency of copulation interruptions, and oviposition success among parental species and hybrid pairs using Fisher Exact tests. We examined the relationship between male abdomen length and cercus size using linear regression. We compared copulation and copulation interruption durations between conspecific and non-conspecific matings using *t*-tests. We compared abdomen length, fecundity, fertility, proportion eggs hatched, developmental timing, and adult lifespan among *E. anna, E. carunculatum,* and hybrids using analysis of variance (ANOVA), after arcsin-transformation of proportion data. When an ANOVA indicated a significant difference existed among the three groups for any measure, we conducted Tukey post-hoc tests to identify the differences among groups. For both forms of premating tactile isolation data, we omitted all cross types with sample size < 6 from statistical analyses. When possible, we combined data (field and lab, or lab generations 1 and 2) to increase statistical power, after confirming with ANOVA that measurements not differ significantly between the two groups. All analyses were conducted in R version 3.1.1 (R Core Team 2015). Means are reported as + 1 SEM.

## Results and Discussion

### Males mate indiscriminately and hybridization occurs at low frequency in nature

At the Whitefish River site, *E. anna* males outnumbered *E. carunculatum* males by a factor of ~1.5. This was observed for both solitary males (*E. anna:* n = 165, *E. carunculatum:* n = 108, over 8 sampling days) and male-female pairs (*E. anna:* n = 44, *E. carunculatum,* n = 28, over 9 sampling days). *E. anna* males attempted tandem with *E. carunculatum* females (46.3%; 19 of 41) as frequently as they did with *E. anna* females (53.7%; 22 of 41; 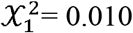, *P* = 0.76; Fig 2). *E. carunculatum* males also attempted tandem with females of both species equally (50.0% (8/16) each; 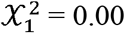, *P* = 1.0; Fig 2). These results show that premating interactions between *E. anna* and *E. carunculatum* are random, similar to observations from other *Enallagma* (Paulson 1974; Fincke et al. 2007; Xu and Fincke 2011) and *Ischnura* species (Sánchez-Guillén et al. 2012; Sánchez-Guillén et al. 2014).

**Fig 2.**
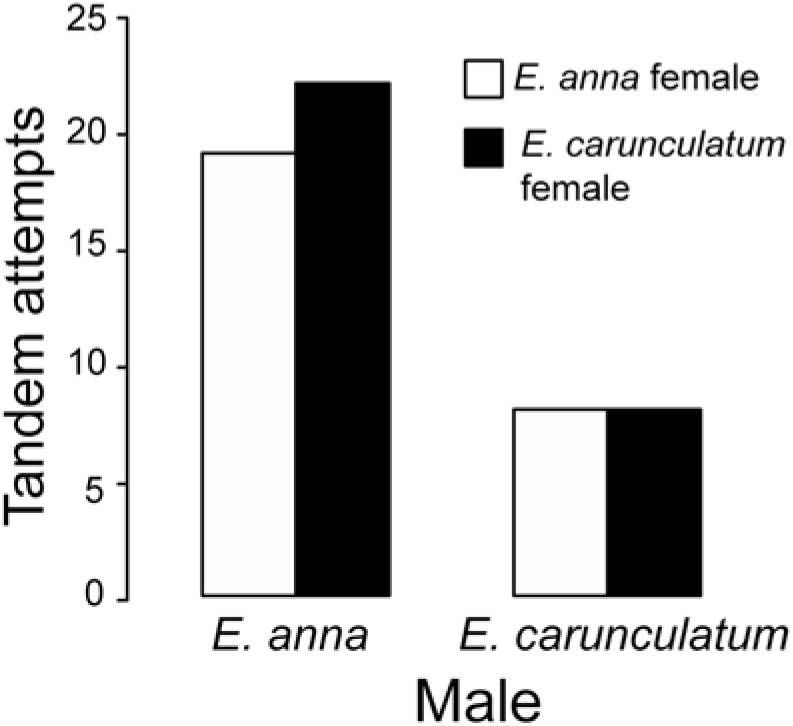
Male visual isolation. Number of male tandem attempts on conspecific and heterospecific females at the Whitefish River site.

Despite this lack of habitat and visual isolation in sympatry, heterospecific pairs were rarely captured in the field. In more than one month at the field site, we captured only two heterospecific male-female pairs, one in each cross direction. Based on the relative frequencies of each pure species, heterospecific pairs occur significantly less often than expected under random mating between *E. anna* and *E. carunculatum* (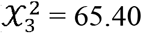, *P* < 1 × 10^−5^). This suggests that although males may frequently initiate tandems with heterospecific females, such pairs likely remain in tandem only briefly. However, even the rare occurrence of both types of heterospecific tandems suggests that pure species may interbreed at low frequencies in the wild, and our collection of field-caught individuals supports this notion: 41 of 630 males and 7 of 547 females we collected possessed intermediate reproductive structure morphologies that were visibly different from either pure species.

### Hybrids are morphologically distinct from either parental species

Among males, the first 5 principal component (PC) scores explained >77% of the cercus shape variance. PC1 (60.05%) distinguished pure species and represented differences in overall cercus length, from short (*E. carunculatum*) to long (*E. anna*), with hybrids showing a range of intermediate scores (Fig 3A). PC2 (7.50%) represented a difference in the relative angles of the upper and lower projections of the cercus, with many hybrids occupying a different space along this axis than parental species. Most field-caught hybrids had distinctly intermediate cercus morphologies, whereas the lab-reared males from heterospecific or backcross pairs possessed morphologies that spanned the entire range of variation between *E. anna* and *E. carunculatum* males (Fig 3A).

**Fig 3.**
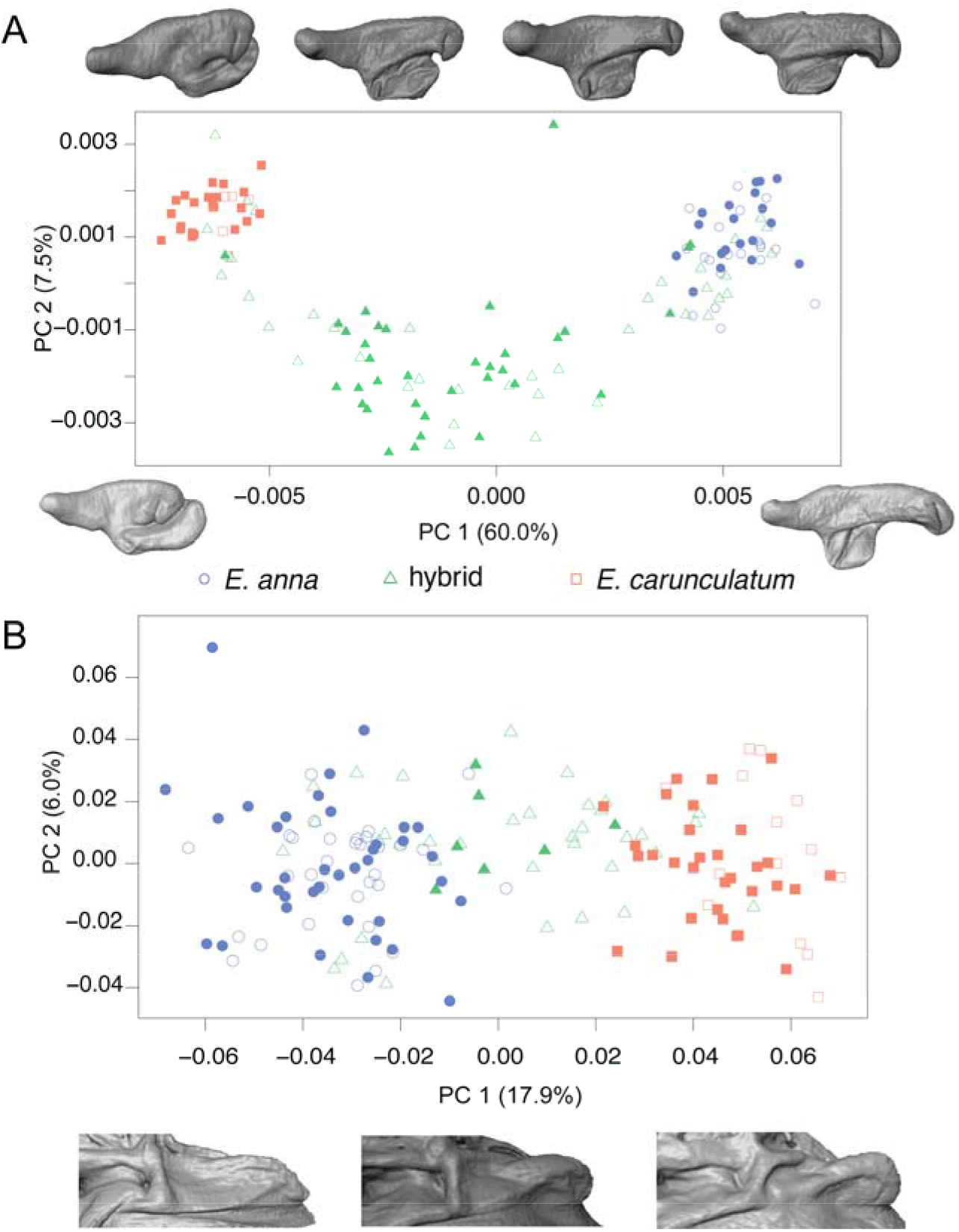
Variation in *E. anna, E. carunculatum,* and hybrid male and female reproductive structure morphologies. **(A)** Distribution of the first two principal components (PC) that represent variation in male cercus shape. Cercus representations above the plot show the range of hybrid male variation across the PC1 axis and representations below the plot show parental species morphologies. **(B)** Distribution of the first two principal components (PC) that represent variation in female mesostigmal plate shape. Examples of representative parental species and hybrid morphologies are shown below the plot (left = *E. anna,* middle = hybrid, right = *E. carunculatum*). The percentage of variation explained by each PC axis is shown in parentheses. Open symbols represent lab-reared individuals, filled symbols represent field-caught individuals.

Among females, the first 6 principal components scores accounted for >42% of the variance in mesostigmal plate shape. *E. anna* and *E. carunculatum* specimens formed separate clusters on PC1 (17.9%), but there was considerable overlap between hybrids and *E. anna* on PC1 (Fig 3B). This overlap might reflect limitations of the resolving power of our morphometric approach to distinguish intraspecific variation from intermediate hybrid morphology of these complex female structures. Additional PC axes indicated that parental species and hybrid plate shapes showed similar levels of variation in several features, including the angle of the plate’s anterior edge relative to the thorax (PC2; 6.0%), curvatures of the plate’s lateral edge (PC3; 5.5%) and plate surface (PC5; 4.6%), and dimensions of the space between the bilateral plates (PC4; 5.3%). Because slight variation in the manual placement of the fixed landmarks on each female has the potential to contribute to this apparent overlap between *E. anna* and the hybrid females, we repeated the entire analysis beginning with placement of landmarks on a subset of 157 plates selected at random. Repeatability was high among landmark coordinates in both sets (*r* > 0.99) and both replicate analyses produced similar results (Fig S3).

Our behavioral, rearing, and morphometric data confirm that individuals with intermediate reproductive structure morphologies are hybrids between *E. anna* and *E. carunculatum* and not a separate species as originally suggested (Miller and Ivie 1995). Interestingly, the collection of lab-reared hybrids (both F_1_ and backcross) included cercus and plate phenotypes not observed in the field-caught samples (Fig 3). Some lab-reared hybrid morphologies were even indistinguishable from those of the parental species, which could be the result of collecting eggs in the field from mated females that may have been storing sperm from previous conspecific matings. Alternatively, some field-caught adult damselflies that we designated as pure species may in fact have been hybrids that maintained “phenotypic integrity” with one parental species despite having highly admixed genomes (Poelstra et al. 2014). Despite this possibility of occasional misidentification, the majority of field-caught individuals we identified as hybrid possess morphologies that fall well outside of the distributions of either pure species. This is particularly true for cercus shape, which has pronounced differences between *E. anna* and *E. carunculatum.*

The distributions of the field-caught versus lab-reared hybrids also show that hybrid genital morphology appears to be under selection in the wild. In particular, the distribution of male morphologies shows that the field-caught hybrids cluster equally distant from the pure species. This result suggests that although interspecific mating occurs in the field, F_1_ hybrids either rarely backcross with parental species or backcross hybrids rarely survive to reproductive age. Our lab-rearing data show that hybrids can in fact backcross with parental species and advanced backcross individuals are viable and fertile (see below). However, future genomic studies will be needed to reveal the direction and genomic extent of introgression and the frequency of F_1_ versus advanced-generation hybrids in the wild.

### Mechanical incompatibilities cause substantial, asymmetric reproductive isolation

Between pure species, precopulatory mechanical RI was incomplete in both directions of interspecific cross, and RI appears asymmetric: 25% (7/28) of *E. anna* males achieved tandems with *E. carunculatum* females, whereas 66.7% (6/9) of *E. carunculatum* males achieved tandems with *E. anna* females (Fig 4A). These data show that mechanical isolation is relatively weak between *E. carunculatum* males and *E. anna* females, which presents the opportunity for interspecific matings. Mechanical isolation due to males’ inability to grasp heterospecific females is frequently evoked as the major contributor to RI in coenagrionid damselflies (Paulson 1974; Robertson and Paterson 1982; Fincke et al. 2007; Bourret et al. 2012; Wellenreuther and Sánchez-Guillén 2016), although several exceptions exist (Paulson 1974; Tennessen 1975; Bick and Bick 1981; Forbes 1991; Miller and Fincke 2004). Our results suggest that mechanical incompatibilities are not sufficiently strong enough to completely exclude the possibility of hybridization in *Enallagma*. Additionally, it has been suggested that species with longer cerci are better at grasping females of other species (Paulson 1974), but our data show that *E. anna* males, whose cerci are roughly twice as long as *E. carunculatum* cerci, were less capable of grasping heterospecific females compared to *E. carunculatum* males.

**Fig 4.**
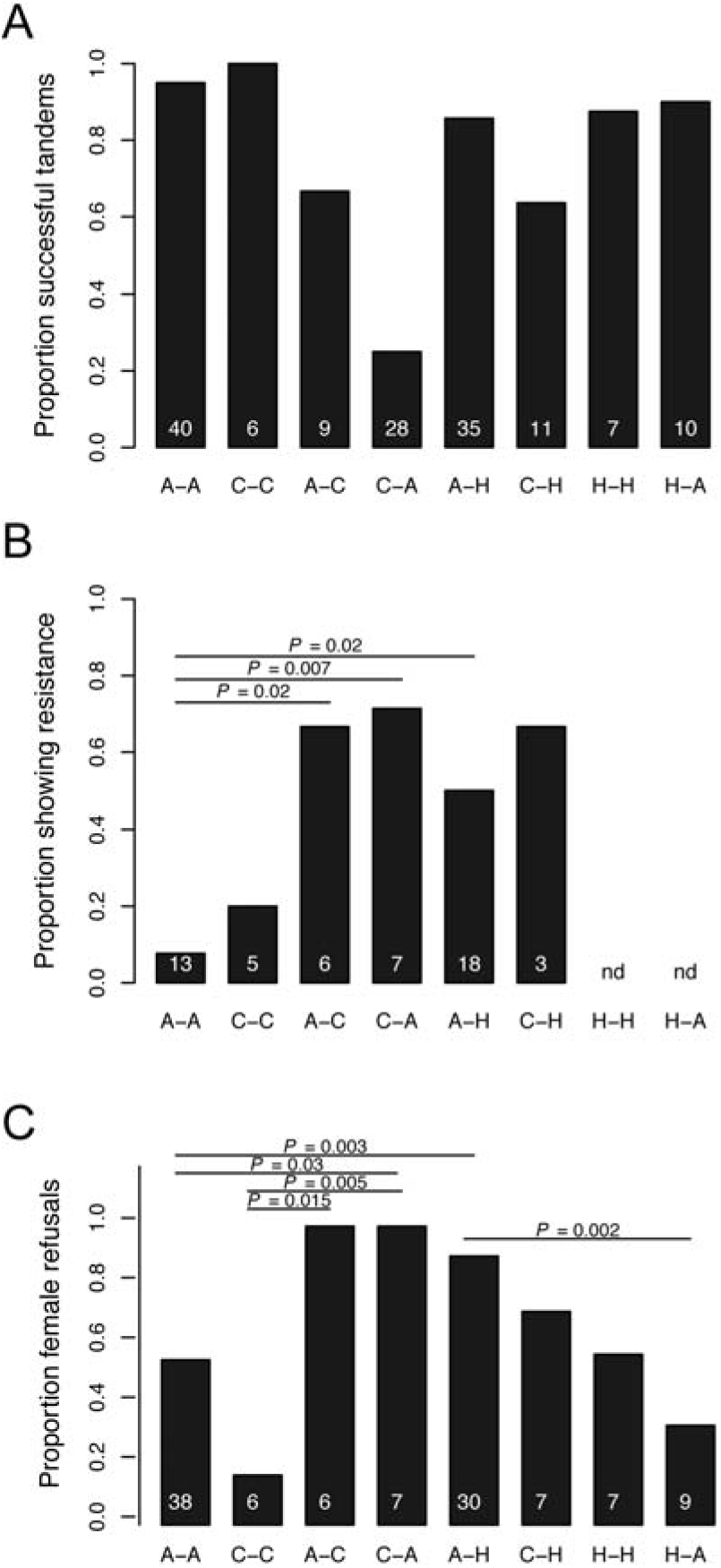
Sequentially-acting mechanisms of prezygotic reproductive isolation. **(A)** Mechanical isolation. **(B)** Proportion of tandems in which females displayed resistance behaviors (field-caught only). **(C)** Proportion of tandems in which females refused to copulate (field-caught and lab-reared data). Crosses shown on the x-axis list female first. A = *E. anna*, C = *E. carunculatum,* H = hybrid. Numbers at the base of the bars in panels A-C show the numbers of male-female pairs that were measured. “nd” refers to cross types for which no data were collected.

The existence of incomplete precopulatory mechanical incompatibilities between *E. anna* and *E. carunculatum* suggests that the intermediate cercus morphology of hybrid males might reduce their ability to form tandems with pure species females. Eighty-six percent (30/35) of hybrid males we tested achieved tandem with *E. anna* females, and 63.6% (7/11) achieved tandem with *E. carunculatum* females (Fig 4A). Thus, male hybrids achieved tandem with both pure species more frequently than males of either pure species achieved tandem with heterospecific females. These results show that although hybrid males were less successful at forming tandems with females than conspecific males, they were more successful than heterospecific males.

Mechanical incompatibility involving the primary genitalia (intromittent organs) may also cause RI. No heterospecific matings occurred during our behavioral observations, so we could not directly measure copulatory mechanical RI between *E. anna* and *E. carunculatum.* However, among the tandem pairs involving hybrids in which the female initiated copulation (2 *E. anna*, 2 *E. carunculatum*, and 3 hybrid females), all 7 pairs achieved genital coupling. Although this sample size is modest, this result suggests that no copulatory mechanical incompatibility exists between hybrids and parental species. This is not unexpected, as *E. anna* and *E. carunculatum* penes have similar morphologies (Kennedy 1919). Taken together, the results from these mating assays show that as the morphological mismatch between interacting male and female mating structures increases, the possibility of forming tandem and mating decreases.

### Tactile incompatibilities cause substantial RI when mechanical isolation is incomplete

A significantly greater proportion of lab-reared *E. anna* females (12/22) engaged in resistance behaviors during conspecific tandems than did field-caught *E. anna* females (1/13, Fisher exact test, *P* = 0.01). For this reason, we analyzed presence/absence of female resistance during tandem separately for field-caught and lab-reared populations. In the field, *E. anna* females were significantly more likely to resist during tandems with heterospecific males (67%; 4 of 6) or hybrid males (50%; 9 of 18) than with conspecific males (7.7%; 1 of 13; Fisher exact tests, *P_heterospecific_* = 0.02, *P_hybrid_* = 0.02; Fig 4B). Additionally, 71.4% (5/7) of *E. carunculatum* females displayed resistance behaviors during tandem with *E. anna* males in the field, which was significantly greater than 7.7% of *E. anna* females (*P* = 0.007; Fig 4B).

Surprisingly, lab-reared *E. anna* females resisted during tandems with conspecific males as frequently as they resisted during tandems with hybrid males (54.5%, 12 of 22 vs. 81.8%, 9 of 11, respectively; *P* = 0.25; Fig S4). *E. anna* and hybrid females also showed similar levels of resistance during tandem with *E. anna* males (14.3%, 1 of 7 of hybrid females resisted; *P* = 0.09; Fig S4). A comparison of the two reciprocal *E. anna* × hybrid crosses, however, showed that *E. anna* females were significantly more likely to resist during tandem with hybrid males (81.8%) than were hybrid females (14.3%; *P* = 0.01; Fig S4). Female resistance during tandem with a conspecific male is not unusual (Tennessen 1975; Fincke 2015), but because the field-caught and lab-reared *E. anna* populations behaved so differently, and the field data reflects behavior in a natural setting, we used the field-caught female data to calculate this form of tactile isolation (Table S1).

Field-caught and lab-reared females showed similar copulatory refusal rates: 94.7% (18/19) field-caught and 81.8% (9/11) lab-reared *E. anna* females refused hybrid males (Fisher exact test, *P* = 0.54), and 69.2% (9/13) field-caught and 51.9% (12/25) lab-reared *E. anna* females refused conspecific males (*P* = 0.31). We therefore pooled field-caught and lab-reared data to analyze female copulation refusal or acceptance. Ninety percent (27/30) of *E. anna* females taken in tandem by hybrid males refused to copulate, which was significantly greater than the 55.3% (21/38) of *E. anna* females that refused conspecific males (P = 0.003; Fig 4C). All six *E. anna* females observed in tandem with *E. carunculatum* males refused to copulate, although this level of refusal was not statistically different from the conspecific refusal rate (*P* = 0.07; Fig 4C). This is likely due to the low number of heterospecific pairs we could observe. *E. carunculatum* females, in contrast, were significantly more likely to refuse an *E. anna* male (100%, 7 of 7) than a conspecific male (16.7%, 1 of 6; *P* = 0.005; Fig 4C). *E. anna* females also refused to mate with *E. carunculatum* males more frequently than did *E. carunculatum* females (*P* = 0.015). We obtained a similar result for the reciprocal cross, where more female *E. carunculatum* females refused *E. anna* males than did *E. anna* females (*P* = 0.03; Fig 4C).

Females’ behavioral responses to different types of males reveal strong assortative mating between *E. anna* and *E. carunculatum* when premating mechanical isolation fails. Tactile isolation also predicts that pure species females should refuse to mate with hybrid males because intermediate cerci fail to relay the proper tactile species recognition signal to the female. Our behavioral data support this prediction for *E. anna* females, which mated with hybrid males less frequently than with conspecific males. The finding that some *E. anna* females mated with hybrid males, but none mated with *E. carunculatum* males suggests that females display some latitude in their preferences and are more likely to refuse males whose cercus morphology greatly deviates from a conspecific phenotype. Although incomplete mechanical isolation has been documented in several *Enallagma* species pairs, few cases of hybridization are known, based on morphological or genetic evidence (Catling 2001; Turgeon et al. 2005; Donnelly 2008). This suggests that even with incomplete mechanical isolation, tactile isolation might prevent interbreeding among most *Enallagma* species. A full understanding of tactile isolation will require quantitative study of the mechanoreceptors on female plates to understand how patterns of phenotypic variation might contribute to RI.

The relative sizes of male and female reproductive structures may influence both mechanical and tactile mechanisms of RI. Larger males tended to have larger cerci, as indicated by regressing cercus volume on abdomen length (*E*. *anna*: *F*_1, 26_ = 18.80, *R*^2^ = 0.397, *P* = 0.0002; *E. carunculatum: F*_1, 17_ = 7.744, *R*^2^ = 0.273, *P* = 0.013). Hybrids, however, showed a weaker relationship between body size and cercus size, because hybrids display more variation in cercus morphology than either parental species (*F*_1,55_ = 6.70, *R*^2^ = 0.092, *P* = 0.01). A size mismatch in male and female structures either within or between species may contribute to mechanical incompatibilities, although our current data do not allow us to examine that relationship robustly.

### Postmating barriers contribute little to reproductive isolation

Compared to the strong premating RI caused by mechanical and tactile incompatibilities of male and female reproductive structures, we found relatively weak RI from postmating barriers. Copulation duration was similar among conspecific mating pairs and pairs including at least one hybrid partner (*t*_25_ = −0.028, *P* = 0.98; Fig 5A). Sixty percent (6/10) of conspecific matings experienced interruptions, which was not significantly different from the hybrid matings (61.5%, 8 of 13; Fisher exact test, *P* = 1.0). The total duration of these interruptions was also not significantly different between conspecific or hybrid pairs (*t*_13.26_= −1.51, *P* = 0.15; Fig 5B). Although it has been suggested that Lepidoptera (Lorkovic 1958) and *Ischnura* (Córdoba-Aguilar and Cordero-Rivera 2008) use copulatory morphology or stimulation to identify conspecifics, our results indicate that this type of tactile discrimination during copula does not occur in *Enallagma*.

**Fig 5.**
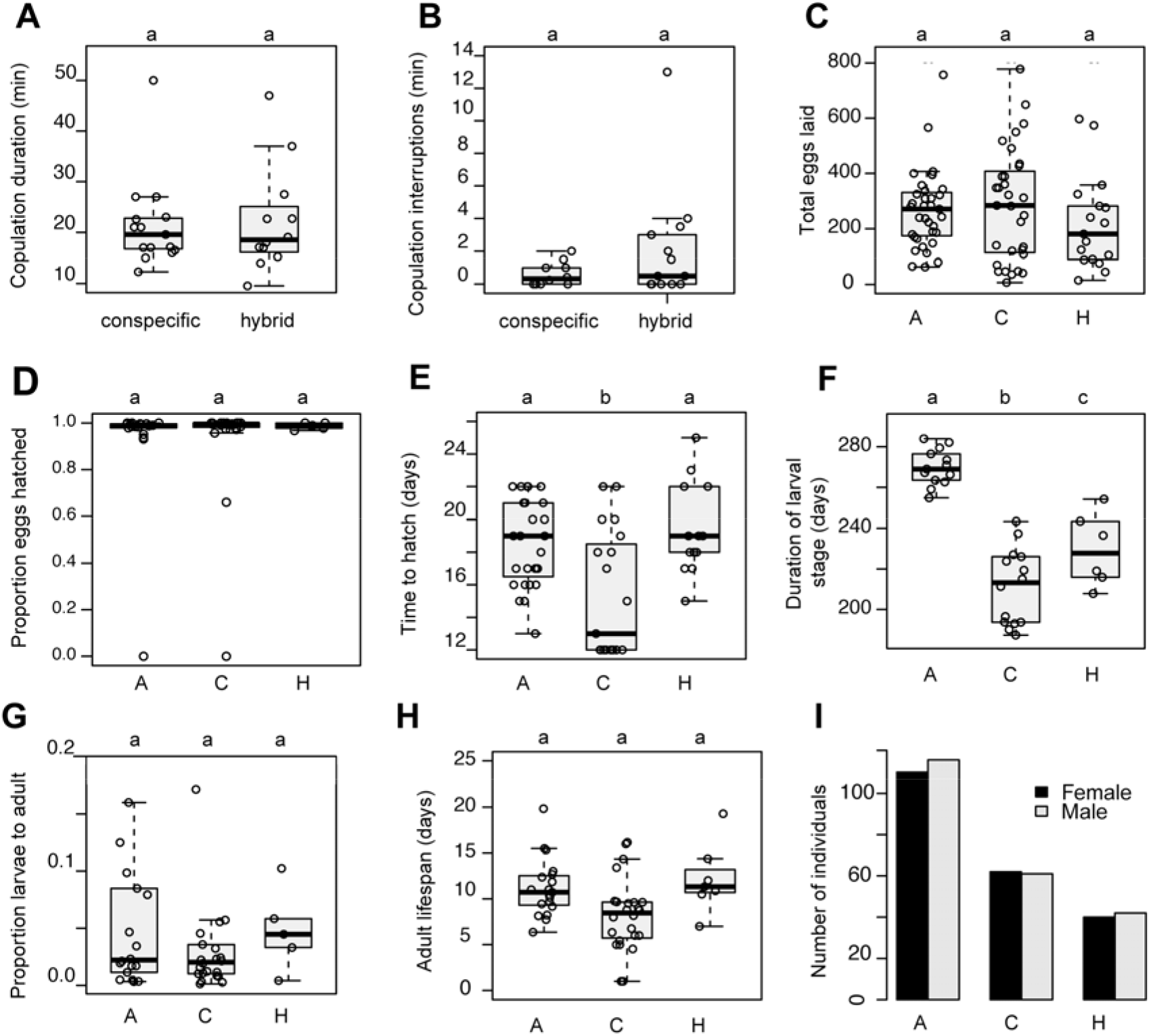
Sequentially-acting mechanisms of postmating reproductive isolation. **(A)** Copulation duration. **(B)** Length of copulation interruptions. **(C)** Fecundity (field-caught and lab-reared data). **(D)** Proportion hatched eggs per clutch. **(E)** Embryonic development timing. **(F)** Larval maturation timing. **(G)** Larval survivorship. **(H)** Adult lifespan. **(I)** Adult sex ratios. In panels A and B, each point represents one male-female pair. In panels C-G, each point represents one clutch. Within each panel, letters indicate homogeneous groups assigned at the statistical cutoff at α = 0.05. Boxplots show the interquartile range. The line within the box shows the median and whiskers extend to the most extreme observation within ±1.5 times the interquartile range. Each open circle represents one mating pair (A, B), one clutch (C-E, G), or the clutch mean (F,H).

Similar proportions of *E. anna* (81.8%; %, 9 of 11) and hybrid females (83.3%, 5 of 6) oviposited after mating with *E. anna* males (Fisher exact test, *P* = 1.0). Two *E. anna* females mated with hybrid males, but neither laid any eggs. In contrast, two *E. carunculatum* females mated with hybrid males and both oviposited. Two of the three hybrid females that mated with hybrid males also oviposited. Dissections of females that failed to oviposit confirmed that they had been inseminated and possessed mature eggs, and dissections of hybrid males in these matings confirmed that hybrid males produce motile sperm. *E. anna, E. carunculatum,* and hybrid parings also produced comparable numbers of eggs (*F*_2,80_ = 0.79, *P* = 0.46; Fig 5C). Although there appears to be a trend towards smaller clutches or complete failure to oviposit in females mated to hybrids, small samples prevent us from drawing strong conclusions about whether tactile incompatibilities might contribute to postcopulatory isolating mechanisms.

The second generation of lab-reared damselflies consisted solely of *E. anna* and advanced generation hybrid clutches, because in generation 1, *E. carunculatum* adults emerged earliest and few were available for crosses with *E. anna* or hybrids. In generation 2, *E. anna* and hybrid clutches had similar fertilization rates (*F*_1, 17_ = 0.51, *P* = 0.49). In generation 1, *E. anna, E. carunculatum,* and hybrid clutches had similar proportions of hatched eggs (Kruskal-Wallis 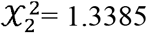, *P* = 0.51; Fig 5C). In generation 2, *E. anna,* and hybrid clutches had similar proportions of hatched eggs (*t*_17.97_= 0.49404, *P* = 0.63, Fig S6). Oviposition date had a significant effect on hatch timing in generation 1 (*F*_1,41_ = 49.1, *P* = 1.6 × 10^−8^), but not in generation 2 (*F*_1,41_ = 2.96, *P* = 0.11). We therefore analyzed hatch timing separately for each generation. In generation 1, *E. carunculatum* larvae hatched earlier (15.4 + 0.9 days, n =19 families) than *E. anna* (19.2 + 0.7 days, n =17 families) and hybrid larvae (20.0 + 1.3 days, n =7 families; ANCOVA with oviposition date as covariate, *F*_2, 39_ = 10.8, *P* = 2 × 10^−4^). In generation 2, *E. anna* and hybrid hatch rates did not differ significantly (*t*_11.92_= −1.22, *P* = 0.25; Fig 5D). If *E. carunculatum* larvae develop at a faster rate in the wild as they did in the lab, this could contribute to RI via seasonal temporal isolation, in which early-emerging *E. carunculatum* adults are less likely to encounter, and thus potentially interbreed with, *E. anna* adults. Detecting and measuring this potential temporal barrier will require regular sampling throughout the breeding season.

An anomalous water quality problem at the Aquatic Research Facility where larvae were housed caused substantial larval mortality of generation 2, so we analyzed larval development timing for generation 1 only (Fig 5F). An ANCOVA with oviposition date as a covariate and Tukey post-hoc tests indicated that hybrids and parental species spent significantly different lengths of time in the larval stage (*F*_2, 29_ = 97.3; *P* < 1.4 × 10^-13^). *E. carunculatum* (n =13 families) larvae reached adulthood an average of 58.6 + 2.5 days earlier than *E. anna* (n =14 families; *P* < 1 × 10^−5^) and 18.2 + 7.3 days earlier than hybrids (n =6 families; *P* = 0.056). Hybrid larvae also developed significantly faster than *E. anna* (*P* = 3 × 10^−5^). Although *E. carunculatum* larvae developed faster than *E. anna* and hybrid larvae in the lab, mean adult abdomen length was similar among all three groups for both males (*F*_2, 19_ = 0.334; *P* = 0.72) and females (*F*_2, 21_ = 3.30; *P* = 0.57; Fig S5). These results suggest that hybrid development was not affected by intrinsic genetic incompatibilities.

Larval survivorship in the lab was similar for both parental species’ and hybrid clutches (Kruskal-Wallis 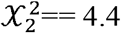, *P* = 0.1; Fig 5G). Of those individuals that reached adulthood, adult lifespans under laboratory conditions did not differ significantly (ANCOVA with emergence date as covariate, *F*_2, 48_ = 1.35, *P* = 0.29; Fig 5H). Of those individuals that reached adulthood, adult lifespans under laboratory conditions did not differ significantly (ANCOVA with emergence date as covariate, *F*_2, 48_ = 1.35, *P* = 0.29; Fig 5H). Finally, adult sex ratios were not significantly different from the expected 1: 1 ratio for any group (Fig 5I), which shows that among pure species and hybrids, both sexes had similar viability. The combination of our postmating isolation results demonstrate that neither strong intrinsic nor extrinsic (*e.g.*, ecological selection against hybrids in the field) postzygotic barriers exist between *E. anna* and *E. carunculatum*.

### Divergent reproductive structures cause reproductive isolation early during speciation

Fig 6 shows the cumulative strength of RI barriers measured for each reciprocal cross. Premating mechanical and tactile incompatibilities form the most substantial barriers to gene flow between *E. anna* and *E. carunculatum,* whereas later-acting barriers contribute little to total RI. Our results thus unequivocally demonstrate the potential of divergent mating structures to cause RI in the early stages of speciation via mechanical and tactile mechanisms. These incompatibilities also appear to provide particularly strong barriers to gene flow, as they act as both a premating barrier between pure species and also as a postzygotic barrier that reduces hybrid male mating success. Such incompatibilities represent a potent barrier to gene flow and may be a common characteristic of traits that are under sexual selection within species (Stratton and Uetz 1986; Naisbit et al. 2001; Höbel et al. 2003; Svedin et al. 2008; Van Der Sluijs et al. 2008).

**Fig 6.**
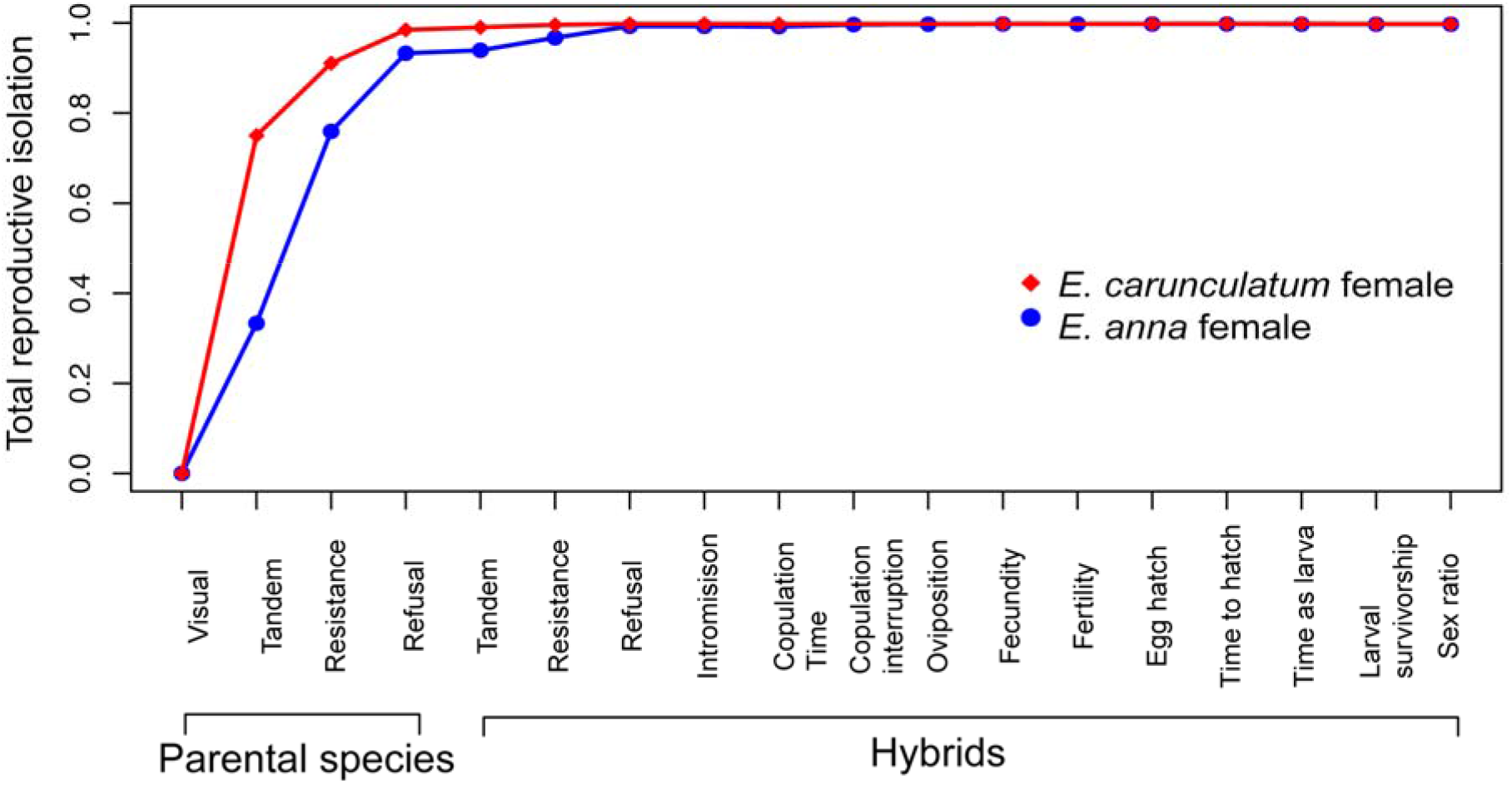
Sequential strength of reproductive isolating barriers, beginning with male-female encounter and proceeding through the reproductive sequence and life history. Estimates of the strength of the first four barriers were obtained from conspecific and heterospecific crosses only, and estimates of the remaining barriers also include crosses involving hybrid individuals. Estimates for the values of the strength of three barriers from the *E. carunculatum* female × hybrid male cross (copulation interruption duration, oviposition, and fertility) are represented by the best-fit line at these barriers.

Our results also show that premating barriers appear to have evolved first in *Enallagma*. Because *E. anna* × *E. carunculatum* hybrids appear to survive as well as parental species and suffer no intrinsic fertility deficits, the primary factor likely to affect their fitness is with whom they can mate. We observed that *E. anna* females often refuse to mate with conspecific males, indicating strong intraspecific discrimination. If the male cerci are under sexual selection similar to non-intromittent mating structures in other taxa (reviewed in Simmons 2014), and if females rely on the same tactile cues for both intraspecific mate choice and species discrimination, then female discrimination among conspecific males could extend to discrimination of heterospecifics. Prezygotic RI has been shown to evolve rapidly under laboratory settings due to assortative mating, independent of local adaptation (Castillo et al. 2015), which supports the plausibility of rapid evolution of RI driven by sexual selection in the wild. Female discrimination against males with intermediate cerci also provides an opportunity for reinforcement to strengthen premating isolation between *E. anna* and *E. carunculatum*— a potential example of sexual selection rather than natural selection driving reinforcement (Naisbit et al. 2001). Reinforcement could result in shifting or narrowing of female preferences (Ritchie 1996) or an increase in female discrimination in regions of sympatry (Noor 1999), two ideas that deserve further study in these species. Alternatively, evolution of cercus morphology may be driven by sexual conflict over mating rate, in which selection favors females that are less easily grasped by males (Fincke et al. 2007).

Many researchers have dismissed genital mechanical incompatibilities as having an important role in RI and speciation (reviewed in Shapiro and Porter 1989; Eberhard 2010), primarily because of the small number of convincing cases that show strict support for it. We might be better equipped to investigate the reproductive consequences of the widespread pattern of rapid, divergent evolution of male genitalia if we broaden our scope to include explicitly tactile mechanisms. This may require dropping the genital “lock-and-key” imagery – which often evokes an “all or nothing” scenario in causing RI – in favor of a framework that allows for more variation, similar to our understanding of auditory, visual, and chemical communication signals. Indeed, our data show that mechanical isolation can be strong yet incomplete, and that tactile isolation can form a strong subsequent mating barrier. A full understanding of the contribution of mechanical incompatibilities in RI will require detailed morphological study to understand how male and female structures interact (Willkommen et al. 2015) and which features cause morphological mismatch. A deeper understanding of tactile RI mechanisms will require detailed studies of sensory mechanisms and the neurobiological basis of female reproductive decisions, all of which are admittedly challenging to investigate. Where females discriminate against heterospecific reproductive structures (*e.g.*, Bath et al. 2012), the female nervous system poses a potentially more complex spectrum of incompatibilities compared to genitalia. Taxa such as damselflies or stick insects (Myers et al. 2016) provide ideal systems to begin to tease apart mechanical and tactile contributions to RI. Neural circuits that integrate olfactory and auditory cues with internal physiological processes to influence female mating decisions are being mapped in *Drosophila* (Bussell et al. 2014; Feng et al. 2014; Zhou et al. 2014), paving the way for similar mechanistic understanding of sensory modalities in emerging model systems. Although odonates have a unique mode of mating that presents multiple opportunities for both mechanical and tactile mismatch, our results highlight the potential contribution of tactile signals involving the genitalia to RI among internally fertilizing animals.

